# Task feedback suggests a post-perceptual component to serial dependence

**DOI:** 10.1101/2022.03.19.484939

**Authors:** Jacqueline M. Fulvio, Bas Rokers, Jason Samaha

## Abstract

Decisions across a range of perceptual tasks are biased toward past stimuli. Such serial dependence is thought to be an adaptive low-level mechanism that promotes perceptual stability across time. However, recent studies suggest post-perceptual mechanisms may also contribute to serially-biased responses, calling into question a single locus of serial dependence and the nature of integration of past and present sensory inputs. We measured serial dependence in the context of a 3D motion perception task where uncertainty in the sensory information varied substantially from trial to trial. We found that serial dependence varied with stimulus properties that impact sensory uncertainty on the current trial. Reduced stimulus contrast was associated with an increased bias toward the previous trial’s stimulus direction. Critically, performance feedback, which reduced sensory uncertainty, abolished serial dependence. These results provide clear evidence for a post-perceptual locus of serial dependence in 3D motion perception and support the role of serial dependence as a response strategy in the face of substantial sensory uncertainty.

## Introduction

In recent years, a number of perceptual phenomena have been explained by appealing to the idea that perceptual reports are not uniquely determined by external stimulus properties but also depend on internal states of the observer. For instance, the success of Bayesian-observer models in accounting for biases in perception support the role of internal states in perception (Knill & Richards, 1996; Weiss et al., 2002; Knill, 2007; Girshick et al., 2011; Rokers et al., 2018). A recent phenomenon that illustrates the contribution of internal state is serial dependence: the finding that recent stimulus history can bias current perceptual reports (Fründ et al., 2014; Fischer & Whitney, 2014).

Serial dependence is thought to be adaptive. Because most features in the environment are stable, especially over short time scales, the brain may incorporate recent information into current estimates as a means of reducing noise in instantaneous sensory signals. Here, we sought to understand the contribution of sequential effects to estimating 3D motion direction and the extent to which such serial biases are impacted by task feedback and sensory uncertainty.

Serial dependencies have been observed across a range of high- and low-level perceptual dimensions, task structures, and sensory modalities (Burr & Cicchini, 2014; Fischer & Whitney, 2014; Liberman et al., 2014; Cicchini et al., 2014; Xia et al., 2016; Alais et al., 2017; Bliss et al., 2017; Cicchini et al., 2017; Cicchini et al., 2018; Kiyonaga et al., 2017; van Bergen & Jehee, 2017; Clifford et al., 2018; Fornaciai & Park, 2018; Liberman et al., 2018; Alexi et al., 2018; Manassi et al., 2018; Suárez-Pinilla et al., 2018; Fritsche & de Lange, 2019; Van der Burg et al., 2019; Pascucci et al., 2019; Samaha et al., 2019; Barbosa & Compte, 2020; Kim et al., 2020; Cicchini et al., 2021; de Azevedo Neto & Bartels, 2021; Ceylan et al., 2021; Kim & Alais, 2021; Pascucci & Plomp, 2021; Manassi & Whitney, 2022; Goettker & Stewart, 2022). It is clear that serial dependence is not simply due to hysteresis in motor processes as the effect does not depend on the need for a motor response (Fisher & Whitney, 2014; Cicchini et al., 2018; Manassi et al., 2018; Fornaciai & Park, 2018). However, beyond that, serial dependencies appear to take on many forms and arise from different sources.

Numerous perceptual bases for serial dependence have been described. Recent work has identified retinal error signals as one source of serial dependence in oculomotor behavior (Goettker & Stewart, 2022), isolating the effects to the earliest stages of visual processing. Serial dependence has been demonstrated to be retinotopic for orientation judgments for stimuli up to 22 degrees of visual angle apart (Collins, 2019), which led to the speculation that serial dependence may also arise at later stages of visual processing. The bias was thought to arise from the activation of inferior temporal neurons with wide receptive fields by orientation-selective neurons in striate and extrastriate cortex, which then feed back to lower visual areas., This feedback was thought to elevate activation of neurons with similar orientation tuning, resulting in the biasing the population response toward the recently encountered orientation on the next trial. Using a reverse correlation technique with classification images, Murai & Whitney (2021) demonstrated that serial dependence biases the priors (‘perceptual templates’) used by the visual system to selectively weigh the visual input according to task demands. Serial dependence has also been shown to operate on perception of features and integrated objects (Collins, 2021), in the integration of dynamic optic flow and static form information in estimates of heading direction (Wang et al., 2022), and to directly alter appearance of perceptual stimuli (Collins, 2020; Cicchini et al., 2017; Cicchini, et al., 2021; Manassi & Whitney, 2022). Other work using orientation judgment tasks has shown attractive serial dependence when past and previous stimuli were close in orientation (Cicchini et al., 2017).

However, using a similar task, attractive serial dependence has also been argued to operate at the decision level (Fritsche et al., 2017; see also Pascucci, et al., 2019), thus implicating post-perceptual contributions. Further evidence for post-perceptual contributions comes from the demonstration of negative (i.e., repulsive) serial dependence (Pascucci et al., 2019) or serial dependence disappearing altogether (Bae & Luck, 2020) when a previous stimulus is encoded but not reported. Moreover, the finding that attractive serial dependence is observed between stimuli that share no overlap in low-level features, but which require the same type of response (Ceylan et al., 2021) supports a post-perceptual component. Recent modeling work has also suggested that serial dependence arises in the mechanisms responsible for the readout of the encoded stimulus in visual cortex (Sheehan & Serences, 2022). Finally, serial dependence has been shown in other cognitive domains including working memory (Fritsche et al., 2017; Bliss et al., 2017; Barbosa et al., 2020) and confidence (Rahnev et al, 2015; Samaha et al., 2019).

The computational principle that determines when and how previous and current inputs are combined also does not appear to be singular. The sensory cue integration literature suggests cues are combined proportional to their associated uncertainty (Trommershäuser et al., 2011; van Bergen & Jehee, 2019). Some accounts of serial dependence are consistent with the cue integration literature and have demonstrated uncertainty-based weighting of previous trial and current trial inputs (Cicchini et al., 2014; Fischer & Whitney, 2014; Cicchini et al., 2018; Samaha et al., 2019; Kim & Alais, 2021). In some cases, serial dependence has been shown to be strongest when past and present trial inputs are similar (Fritsche, 2016; Cicchini et al., 2018; Fritsche & de Lange, 2019; Lidström, 2019). Recent work have also suggested that uncertainty-weighting is not a general principle in that the uncertainty associated with only the current, but not the previous trial, may be incorporated (Ceylan et al., 2021; Gallagher & Benton, 2022). More generally, it has been suggested that serial dependence reflects changes in criteria, resulting from integration with perceptual expectations, as a distinct mechanism from priming (Galluzzi et al., 2022). Finally, it is typically difficult to numerically assess an observer’s estimated reliability of both previous and current trial information, making uncertainty-weighted predictions challenging to test.

In the current study, we investigated serial dependence in the context of 3D motion perception. Although the perception of 3D motion is critical to navigate our environment and interact with our surroundings, the perception of 3D motion in the laboratory is often systematically biased (e.g., Fulvio et al., 2015; Harris & Dean, 2003; Harris & Drga, 2005; Lages, 2006; Rushton & Duke, 2007; Welchman et al., 2004; Welchman et al., 2008). Specifically, 3D motion stimuli tend to be associated with highly variable, but well-characterized, estimates of perceptual uncertainty. For example, observers frequently misreport the motion-in-depth direction of an object such that when approaching, it is reported as receding, and vice versa (Fulvio et al., 2015; Fulvio & Rokers, 2017). Such errors are not simply the result of guessing, but rather arise in a systematic and principled way. First, the frequency with which they occur increases with increased uncertainty associated with the motion signal, as occurs, for example, when the contrast of the stimulus is reduced (Fulvio et al., 2015). Second, these errors are specific to motion-in-depth as they rarely occur for lateral (i.e., leftward and rightward) object motion. Third, these phenomena can be explained by a Bayesian-observer model that optimally weighs incoming sensory information with prior knowledge about object motion in general (Rokers et al., 2018). Finally, the frequency of perceptual misreports decreases with task feedback (Fulvio & Rokers, 2017).

Because of the considerable variability in sensory uncertainty of 3D motion stimuli, and their lawful relationship to stimulus contrast and motion direction, they are an ideal candidate to investigate response biases due to serial dependence. Specifically, we have shown previously that report feedback reduces the motion-in-depth misreports and improves overall performance (i.e., percentage of interceptions). Rapid learning occurs early in the first block of training followed by a much slower improvement over time (Fulvio & Rokers, 2017). Because the performance improvements occur much more rapidly than would be expected based on perceptual learning, we believe the performance improvements are due to the recruitment of the available visual cues in the stimulus display rather than changes in low level sensory processing *per se* (see also Fulvio, Ji, et al., 2020). That is, experience with the 3D display appears to overcome ‘flatness’ priors developed through use of other displays outside the laboratory, where binocular depth cues are typically unavailable. Thus, if serial dependence is diminished with task feedback, the results would support a post-perceptual component of serial bias in this task. Thus, by exploiting the relationship between 3D motion stimuli and their associated perceptual uncertainty, we aimed to gain insight into the computational principles underlying serial dependence and identify the source of serial effects.

## Methods

### Participants

Data from 72 participants were used in the current analysis. All were recruited from the University of Wisconsin - Madison undergraduate student population and were naive to the purposes of the study. The study was approved by the University of Wisconsin - Madison institutional review board. All participants gave informed consent and received course credit for their participation. After initial data analysis, data from one participant were excluded from further analysis due to excessive serial bias beyond three standard deviations of the group mean. This resulted in a final set of data from 71 participants (35 females) upon which all results reported in the text are based.

### Experimental design

All participants completed three blocks of a 3D motion-in-depth extrapolation task while wearing a virtual reality head-mounted display. We designed the experiment to resemble the classic video game Pong to make the task intuitive to participants. A major difference between our experiments and the classic video game however, is that we used virtual reality to present stimuli in three dimensions (primarily the x-z plane), whereas classic Pong presents stimuli in 2D (the x-y plane). See Supplementary Materials for a video illustrating the stimulus and experimental procedure. In all, 37 participants from the final set of 71 used in the current analyses completed the task without feedback on the accuracy of their reports, 10 participants completed the task with auditory feedback, and 24 participants completed the task with visual and auditory feedback.

### Visual stimuli

On each trial, the participant fixated the center of a gray circular aperture (7.5 deg radius) within a 1/f noise-mapped planar surface rendered at a viewing distance of 45 cm. A white spherical target (ball, 0.25 deg radius) appeared at fixation and traveled along a random trajectory in the x-z plane for 1 s before disappearing (see **Figure 1**). The target was rendered under perspective projection, so that both monocular (looming) and binocular (disparity) cues to motion-in-depth were present. During the response phase, a rectangular block (paddle, 0.5□cm□×□1□cm□×□0.5□cm) textured with 1/f random noise was presented.

**Fig. 1.**
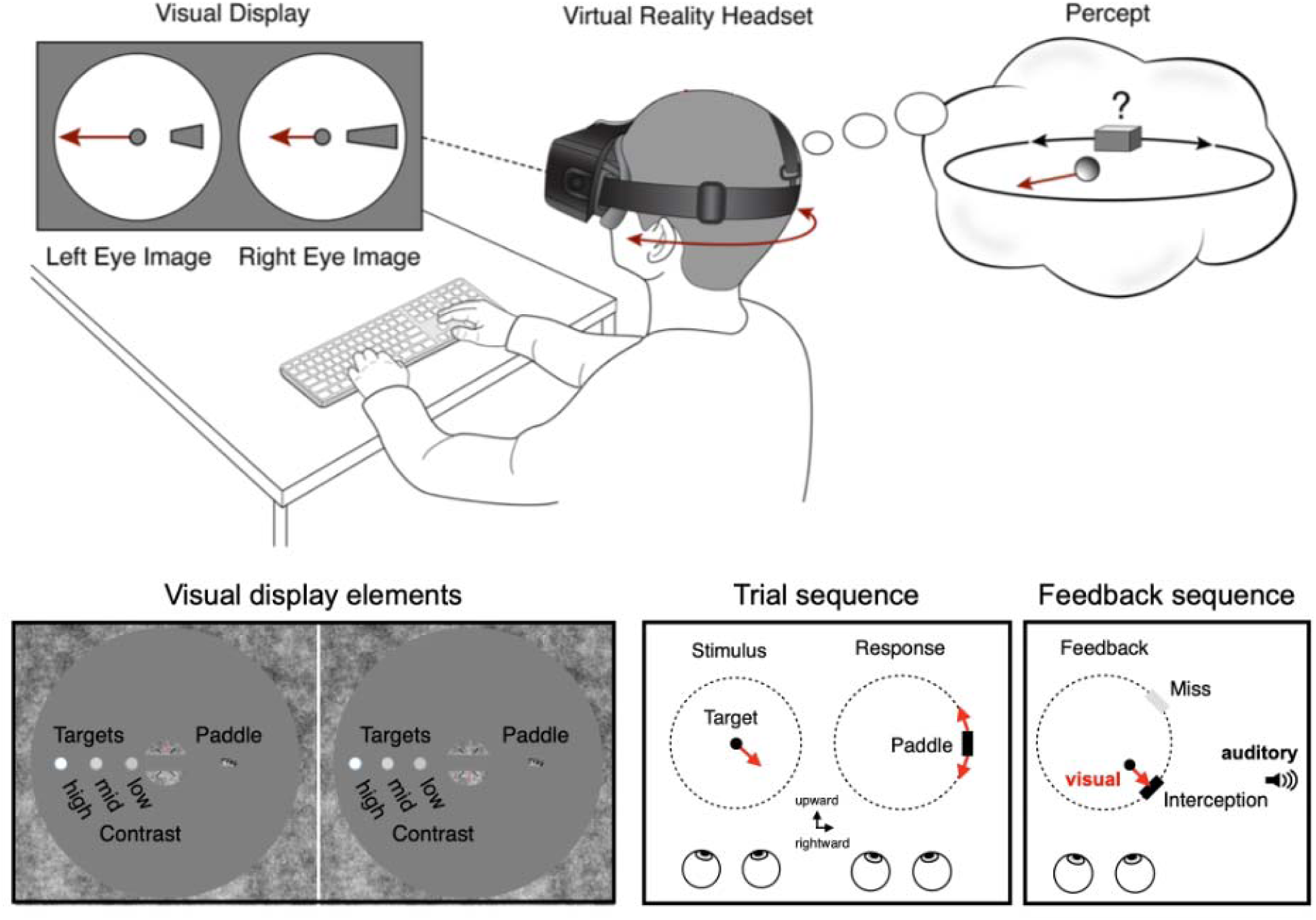
Experimental paradigm. Upper panel: Participants viewed a 3D scene inside a virtual reality headset that consisted of a 1/f noise-mapped planar surface (“wall”) with a circular aperture. Lower panel: A small fixation patch with superimposed nonius lines resided in the center of the aperture. The trial sequence consisted of a white circular target of variable contrast (‘low’, ‘mid’, or ‘high’) appearing at fixation and traversing a random trajectory along the x (lateral) and z (depth) planes for 1 second before disappearing. (Note that for display purposes, the targets are shown separately at eccentricity in the “Visual display elements” panel.) When the target disappeared, participants adjusted the position of a 3D rectangular block (“paddle”) to the location along an invisible orbit (drawn as a dashed circle here for display purposes) that would intercept the target had it continued along its trajectory. One group of participants did not receive feedback and pressed a key to begin the next trial with a new target. A second group of participants received auditory feedback on every trial, and a third group received visual and auditory feedback on every trial. The visual feedback sequence comprised the target reappearing at its last visible location and continuing along its trajectory to the invisible orbit. An interception occurred when the target reached the paddle; otherwise, the setting resulted in a miss. The auditory feedback consisted of an auditory signal corresponding to a “hit” sounded upon interception, otherwise, a “miss” sound was played.

### Procedure

Participants were asked to report the motion direction of the white circular target, i.e. a ball. To do so, participants adjusted the position of the paddle so that it would intercept the target. Paddle position was restricted to an invisible circular orbit centered on the fixation point (**Fig. 1**) and was initialized at a random position on each trial. In the no-feedback group, participants locked in their responses and continued to the next trial after a 1 second fixation interval. In the feedback group, participants instead saw the target reappear at its last location and continue along its trajectory until reaching the (invisible) circular orbit. If the paddle had been placed such that the target was intercepted, a cowbell sound played, otherwise, a “swish” sound played. The next trial then began after the 1-second fixation interval.

Within a block, white target stimuli were presented at three different luminance levels defined in terms of their Weber contrast levels computed by

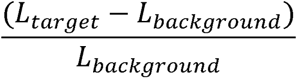

Where *L_target_* and *L_background_* correspond to the luminances of the target and background, respectively. The three target luminance levels were 77.92 cd/m^2^, 69.02 cd/m^2^, 65.68 cd/m^2^ and presented against a gray background (44.56 cd/m^2^), which corresponded to 0.75 (‘high’), 0.55 (‘mid’), and 0.47 (‘low’) Weber contrast levels, respectively. Henceforth we use the labels ‘high’, ‘mid’, and ‘low’ to refer to these contrast levels just as a matter of convenience. The different target contrast levels were presented in a random, counterbalanced order. Across blocks, three rendering conditions were tested in a random order across participants: (i) ‘fixed’ in which the scene did not update according to the participant’s head motion; (ii) ‘active’ in which the scene updated according to the participant’s head motion; (iii) ‘lagged’ in which the scene updated according to the participant’s head motion, but with a random lag of 50 ms on each trial (no-feedback group) or a random lag on each trial drawn from the uniform distribution [0 500 ms] (feedback group). We note that the three rendering conditions were included in the study design to address questions beyond the scope of this investigation and do not materially impact the serial dependence results reported here. These details are described more extensively in a separate study investigating the role of response feedback in the recruitment of visual cues in virtual reality (Fulvio & Rokers, 2017). Participants in the no-feedback and auditory feedback groups completed 225 trials per block for a total of 675 trials. Participants in the auditory + visual feedback group completed 120 trials per block for a total of 360 trials.

### Analysis of current trial performance

Response error on each trial was computed as the difference between presented and reported motion direction. We computed the circular distance between the midpoint of the participant’s paddle setting and the endpoint of the presented target trajectory using the *circ_dist.m* function of the Circular Statistics Toolbox for Matlab (Berens, 2009; **Fig. 2a**). Error distributions were plotted and summarized according to the mean standard deviation of the signed errors, mean absolute error, and proportion of motion-in-depth misreports for comparisons (i). across target contrast level conditions within each feedback condition using one-factor repeated-measures ANOVA. In some cases, the assumption of sphericity was violated, so a Greenhouse-Geisser correction was included in the model;, and (ii). across feedback conditions using two-sample *t*-tests. In some cases, a two-sample *F*-test for equal variances produced significant results, indicating that the two samples violated the equal variance assumption. In those cases, a Welch’s two-sample *t*-test was used where indicated.

**Fig. 2.**
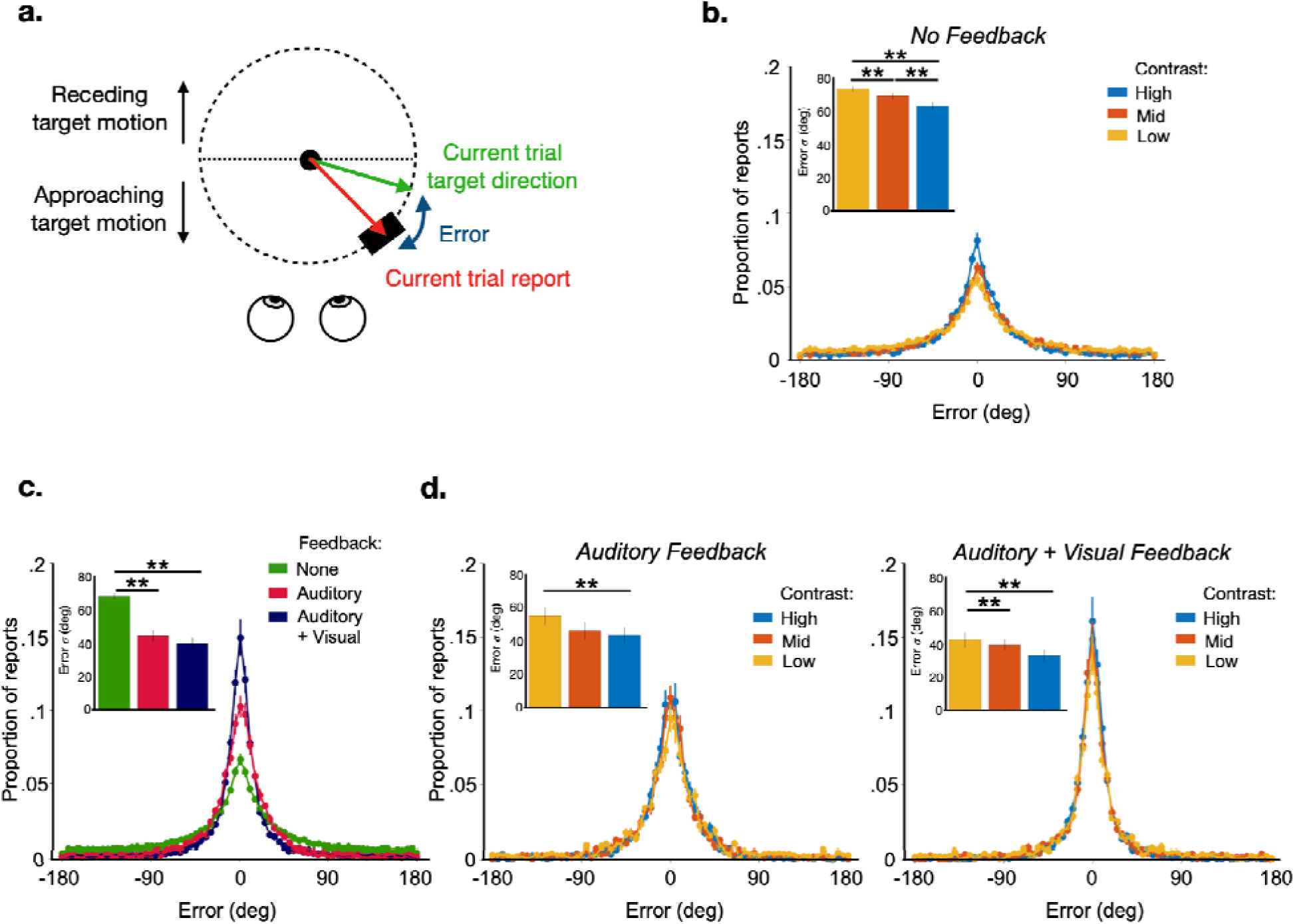
Behavioral performance on the current trial. (**a.**) Error in report on the current trial is defined as the circular distance in degrees between the current trial target direction and the current trial reported direction. (**b.**) Between-subjects error histograms for the group that did not receive feedback for the high (light blue), mid (red), and low (yellow orange) target contrast levels. Left inset depicts the standard deviation of the errors for the three target contrast levels. (**c.**) Between-subjects error histograms for the group that did not receive feedback (green), the group that received auditory feedback (maroon) and the group that received both visual and auditory feedback (dark blue). The inset depicts the standard deviation of the errors for the feedback groups. (**d.**) Same format as (**b.**) for the auditory feedback group (left) and the auditory + visual feedback group (right). Error bars correspond to +/− 1 *SEM*; ** corresponds to significance at the Bonferroni-corrected alpha value = .0167.

### Serial dependence analysis

Serial dependence was quantified using a pipeline similar to that used by Samaha et al. (2019). For each subject, the response error on each trial was computed as the angular distance between the presented and reported motion trajectory. Following previous serial dependence work (Bliss et al., 2017; Fritsche et al., 2017; Samaha et al., 2019), high-error trials were omitted from analysis. In this task, high-error trials were defined as those in which participants misreported the direction of the target’s motion in depth (i.e., when participants reported an approaching target as receding and vice versa, henceforth named “depth misreport trials” contrasted with “correct depth trials”). Such depth misreport trials were more prevalent in the no-feedback condition compared to the feedback conditions (38.3% vs. 18.1% (auditory) and 18.5% (auditory + visual), respectively, on average across conditions). We emphasize that these depth misreports were not simply random settings - the lateral (x) component of the target’s motion was almost always reported correctly on these trials. We also carried out the analyses with depth misreport trials included and include them where appropriate, primarily in Supplemental Materials. To remove any global directional biases for individual participants, we subtracted their mean signed error across trials from each trial. On average, these mean signed errors computed across all trials were small (*M_no-feedback_ =* 0.353°, *SD_no-feedback_ =* 3.173°; *M_auditoryfeedback_=* −0.179°, *SD_auditoryfeedback_ =* 2.622°; *M_auditory+visualfeedback_=* 0.165° *SD_auditory+visualfeedback_ =* 3.391°).

Model-based serial dependence was then quantified using the standard approach of sorting each participant’s demeaned errors according to the angular difference between the presented target direction on the current and previous trials. We note that because the presented target directions spanned the full 360° space, the angular difference scale ranged from [-180°, 180°], twice the range of studies using orientation stimuli. The first trial of each block was omitted because it did not have a previous trial. When trials were removed due to response misreports as described above, the most immediately preceding non-misreport trial served as the previous trial. This approach is reasonable given previous work showing that serial dependence may extend to at least three trials back (Fischer & Whitney, 2014).

The sorted errors for each participant were then smoothed by a 45-degree wide median filter in 1 degree increments. Similar results were obtained with smaller filters. We averaged the smoothed data within each group and fitted the averages of each group with a derivative-of-Gaussian (DoG) function of the following form:

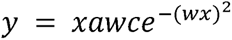

where *x* is the relative orientation of the target of the previous trial, *a* is the amplitude of the peaks of the curve, *w* scales the width of the Gaussian derivative, and *c* is a constant √2/e^-0.5^, which scales the amplitude parameter to numerically match the height of the curve in degrees. Fitting was achieved by minimizing the sum of squared errors using the *lsqcurvefit* MATLAB routine, with amplitude *a* and width *w* parameters free to vary between [-10 deg, 10 deg] and [0.001, 0.5], respectively.

Statistical significance of the group-level DoG fits was assessed through a bias-corrected and accelerated (BCa) bootstrapping procedure. On each of 100,000 iterations, we sampled subjects with replacement and fit a DoG to the average of the bootstrap sampled data. The value of the amplitude parameter was retained after each iteration, resulting in a distribution of the amplitude parameter of our sample. We also computed the jackknife distribution of amplitude parameters by computing all of the leave-one-sample-out DoG fits. The confidence interval and *p*-value of the amplitudes of each group fit when compared to zero (i.e., no serial dependence) were computed using the *BCa_bootstrap* MATLAB function (Van Snellenberg, 2018) which utilized both the bootstrapped and jackknife amplitude distributions.

Between-group comparisons of DoG fit amplitudes were carried out using permutation testing. On each of 100,000 iterations, we shuffled the group labels (e.g., ‘no feedback’ and ‘auditory feedback’) and then assigned each data set with one of the shuffled labels. We then grouped the data by the shuffled labels and fit a DoG to the average of each shuffled group. The value of the amplitude parameter was retained after each iteration, resulting in a permutation distribution of the amplitude parameter. The *p*-value of the original difference in amplitudes between the two groups was derived from the permutation distribution as the proportion of the permutations greater than the original difference.

Following Samaha et al. (2019), we additionally carried out a model-free analysis to verify that the results were not specific to the model-fitting. For each participant, we computed the median (signed) error across trials where the relative difference between the current and previous stimulus fell within the 0-180 deg interval, and subtracted that from the median error on trials within the −180-0 deg interval. Thus, positive and negative values indicate an attractive or repulsive bias, respectively. Statistical testing was performed using two-tailed *t*-tests, one-way ANOVAs, or a linear mixed effects model as described in the main text.

Both model-based and model-free analyses were carried out multiple times on trial sortings according to several questions addressed below. For the basic comparisons between serial dependence on previous target direction, and between performance with and without feedback, trials across all three blocks were used. For block-based comparisons, serial dependence was estimated separately for each of the three blocks. For target contrast-based comparisons, serial dependence was estimated for each of the three target contrast levels, combining trials across blocks.

## Results

### Performance improves with both task feedback and target contrast

We first considered performance as a function of current task conditions. For each trial, we computed the response error as the circular distance between the presented and reported target trajectory (**Fig. 2a**). When feedback was not provided, mean absolute error in motion direction report changed as a function of target contrast (low: M = 55.1 deg (SD = 11.8 deg); mid: M = 51.2 deg (SD = 12.6 deg); high: M = 44.7 deg (SD = 13.2 deg); *F*(1.664,59.905) = 27.310, *p* < .001) as did the proportion of motion-in-depth misreports (*F*(1.699,61.156) = 14.172, *p* < .001) and the standard deviations of the response errors (**Fig. 2b bar graph inset**).

When feedback was provided, mean absolute error in motion direction report was significantly reduced when compared to the performance of the no-feedback group for both feedback types (auditory: M = 28.5 deg (SD = 8.5 deg); *t*(45) = 5.549, *p* < .001; auditory + visual: M = 24.9 deg (SD = 13.7 deg); *t*(59) = 7.702, *p* < .001) with no difference between the two feedback conditions (*t*(32) = 0.7691, *p* .448) with the same pattern observed in the standard deviations of the response errors (**Fig. 2c bar graph inset)**. Relatedly, feedback was associated with a reduction in motion-in-depth misreports (38.3% (no feedback) vs. 18.1% (auditory feedback; *t*(45) = 7.056, *p* < .001) and 18.5% (visual + auditory feedback; *t*(59) = 7.5584, *p* < .001), on average, with no difference between the two feedback conditions *t*(32) = 0.1027, *p* = .919).

Mean absolute error in motion direction report for the auditory feedback group changed as a function of target contrast (low: M = 36.8 deg (SD = 14.9 deg); mid: M = 30.2 deg (SD = 13.2 deg); high: M = 27.8 deg (SD = 10.9 deg); *F*(2,18) = 11.115, *p* < .001) as did the proportion of motion-in-depth misreports (*F*(2,18) = 10.828, *p* < .001) and the standard deviations of the response errors (**Fig. 2d bar graph inset**). Mean absolute error in motion direction report for the auditory + visual feedback group also changed as a function of target contrast (low: M = 28.2 deg (SD = 17.7 deg); mid: M = 25.6 deg (SD = 14.1 deg); high: M = 21.4 deg (SD = 13.2 deg); *F*(2,46) = 9.037, *p* < .001) as did the proportion of motion-in-depth misreports (*F*(2,46) = 3.720, *p* = .032) and the standard deviations of the response errors (**Fig. 2d bar graph inset**). In summary, response errors in the judgment of 3D motion direction decreased within the same observer as stimulus contrast increased. Similarly, response errors decreased when observers received task feedback, compared to when observers did not receive feedback.

We next describe the results of the serial dependence analyses in turn. For completeness, Table 1 at the end of the Results section summarizes all key findings.

**Table 1.** Summary of serial bias results. The first three lines of each cell report the bias obtained in each of the three conditions. Additional lines report significant comparisons among feedback conditions. *NF* = no feedback; *AF =* auditory feedback; *AV* = auditory + visual feedback. Bolded values highlight repulsive biases. *corresponds to *p* < .05; **corresponds to *p* < .01.

| Analysis | Serial bias measure |  |
| --- | --- | --- |
|  | DoG amplitude | Model-free |
| Overall | NF: 2.56° **<br>AF: 0.12°<br>AV: 0.81°<br>-----<br>AF vs. NF * | NF: 3.01° **<br>AF: **-1.01°** *<br>AV: 1.26° *<br>-----<br>AF vs. NF **<br>AV vs. NF *<br>AF vs. AV * |
| By block<br>(B1 / B2 / B3) | NF: 3.58° ** / 2.21° * / 2.58° *<br>AF: 1.21° / 0.05° / **-0.44°**<br>AV: 2.09° / 1.24° / 1.26°<br>-----<br>B3: AF vs. AV * | NF: 4.20° * / 2.90° ** / 2.20° *<br>AF: 1.03° / **-1.99°** * / **-1.24°**<br>AV: 2.80° * / 1.42° / 0.92°<br>-----<br>B2: AF vs. NF **<br>B2: AF vs. AV ** |
| Target contrast -<br>current trial<br>(Low / Mid / High) | NF: 4.37° * / 4.30° ** / 0.17°<br>AF: 1.37° / **-1.49°** / 1.44°<br>AV: 3.28° ** / 0.99° / **-0.98°**<br>-----<br>Mid: AF vs. NF **<br>Mid: AV vs. NF ** | NF: 4.27° * / 3.47° ** / 1.21°<br>AF: **-1.18°** / **-1.90°** * / 0.88°<br>AV: 2.41° ** / 1.14° / 0.04°<br>-----<br>Low: AF vs. NF *<br>Low: AF vs. AV *<br>Mid: AF vs. NF *<br>Mid: AF vs. AV * |
| Target contrast<br>(previous trial)<br>(Low / Mid / High) | NF: 4.21° ** / 3.77° ** / 0.60°<br>AF: 1.61° / **-1.12°** / 0.17°<br>AV: 0.71° / 0.91° / 0.79°<br>-----<br>Mid: AF vs. NF **<br>Mid: AV vs. NF ** | NF: 5.24° ** / 4.70° ** / **-0.70°**<br>AF: 0.64° / **-1.87°** ** / **-1.53°** *<br>AV: 1.42° / 1.84° * / 0.85°<br>-----<br>Low: AV vs NF **<br>Mid: AF vs. NF **<br>Mid: AV vs. AF *<br>High: AV vs. AF * |
| Target contrast<br>(relative current<br>and previous trial)<br>(Curr lower / Curr<br>higher / Curr same) | NF: 1.88° / 1.44° / 5.40° **<br>AF: **-0.002°** / 0.39° / 0.29°<br>AV: 1.26° / **-0.79°** / 1.34°<br>-----<br>Curr same: AF vs. NF **<br>Curr same: AV vs. NF ** | NF: 0.75° / 1.91° / 6.08° **<br>AF: **-1.76°** / **-0.28°** / **-0.84°**<br>AV: 1.20° / **-0.02°** * / 2.20°<br>-----<br>Curr lower: AV vs AF *<br>Curr same: AF vs. NF ** |
|  |  | Curr same: AV vs. NF * |

### Prior stimuli bias current responses

We next computed serial dependence in the 3D motion perception task in the absence of task feedback. In keeping with previous practice, we omitted high error trials (i.e., motion-in-depth misreport trials; but see **Figs. S1-S2** in *Supplementary Materials* for results including all trials). Consistent with previous studies, current responses were biased toward the previous trial’s presented direction (serial dependence amplitude, *a* = 2.56°, 95% CI: [1.39°, 3.87°], *p* < .001) ; **Fig. 3a**). The significant serial dependence was confirmed with our model-free measure of serial bias (*M* = 3.01°; *t*(36) = 3.3050, *p =* .002; **Fig. 3b**). Indeed, the magnitude of bias is consistent with prior work using motion and orientation stimuli (Alais et al., 2017; Fritsche & de Lange, 2019; Samaha et al., 2019; Kim et al., 2020; Celyan et al., 2021; but see Fischer & Whitney, 2014 who reported biases >8° in several conditions).

**Fig. 3.**
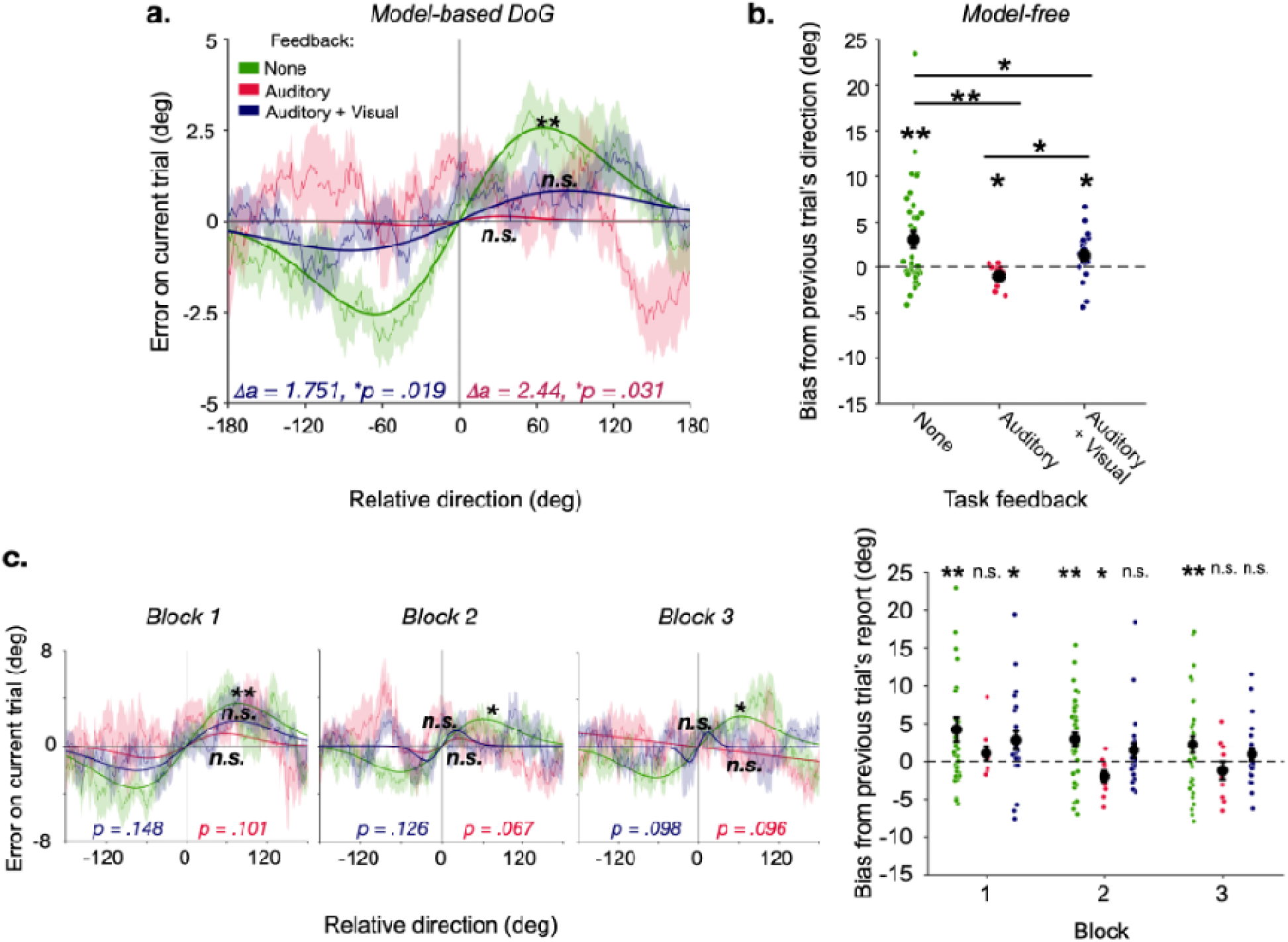
Task feedback reduces serial dependence on previous target direction. **(a.)** Serial dependence curves and DoG fits to error data with high error trials omitted (i.e., on correct depth report trials following correct depth report trials), when no feedback was provided (green), when auditory feedback was provided (red), and when auditory + visual feedback was provided (blue), sorted according to the relative direction of the current trial’s target with respect to the previous trial’s target direction. Shaded bands represent +/-1 *SEM*. **corresponds to bootstrapped *p*-values < .001; n.s. = non-significant; LJ*a* refers to the difference in amplitude parameter between no feedback and auditory feedback groups (red font) and between no feedback and auditory + visual feedback groups (blue font) with corresponding *p*-values derived from permutation testing. **(b.)** Model-free serial bias with respect to the previous trial’s target direction for the three feedback conditions. Circular symbols correspond to individual subject biases; black symbols correspond to group means. Error bars represent +/-1 *SEM*. **corresponds to *p*-values < .01; *corresponds to *p*-values < .05. **(c.)** Same formats as **(a.)** & **(b.)** split by block. One outlier with bias > 45° in the no feedback condition of Block 1 was excluded from the plot.

### Task feedback reduces serial dependence

The significant serial bias reported above extends the finding of serial dependence to the 3D motion domain. To further examine the nature of serial dependence in this task, we analyzed the impact of task feedback. In previous work, we have shown that feedback is associated with an improved ability to intercept the target (Fulvio & Rokers, 2017). Furthermore, our previous work suggests that visual and auditory report feedback improve performance by encouraging the appropriate recruitment of available cues in estimating the motion-in-depth direction (Fulvio & Rokers, 2017; Fulvio, Ji, et al., 2020). This, in turn, reduces the prevalence of motion-in-depth misreports. Our interpretation of the mechanism through which feedback impacts performance in this task is consistent with a post-perceptual locus of feedback-based effects - rather than feedback impacting the quality of early sensory representations, feedback appears to affect the integration of the sensory estimates based on the various cues available in the stimulus. We therefore hypothesized that feedback would also be associated with a reduction in serial dependence as cue recruitment provides more reliable sensory signals.

Consistent with this expectation, task feedback eliminated serial dependence of the current trial’s report on the previous trial’s target motion direction according to the model-based analysis (auditory feedback: *a* = 0.12°, 95% CI: [-2.97°, 1.18°], *p* = .96; auditory + visual feedback: *a* = 0.81°, 95% CI: [-0.39°, 1.45°], *p* = .23; **Fig. 3a**), with no difference in serial dependence observed between the two feedback groups with the model-based measure (lJ*a* = 0.6898°, *p* = .173). However, the model-free analysis suggests a small bias remained with feedback. For the auditory + visual feedback group, this bias remained attractive toward the previous report (*M* = 1.26°; *t*(23) = 2.5225, *p* = .02), whereas the bias was repulsive from the previous report for the auditory feedback group (*M* = −1.01°; *t*(9) = −2.3571, *p* = .043; **Fig. 3b**). The difference in the model-free measure of serial bias for the two feedback groups was significant (*t*(32) = 2.7509, *p* = .01).

Critically, however, both measures revealed a significant reduction in serial dependence with feedback compared to performance when feedback was not provided (auditory feedback: (model-based) lJ*a* = 2.44°, *p* = .031; (model-free): Welch’s two-sample *t*(44.91) = 3.9952, *p <* .001; auditory + visual feedback: (model-based): lJ*a* = 1.75°, *p* = .019; (model-free): Welch’s two-sample *t*(53.36) = 1.6845, *p =* .049).

Investigating how serial dependence changes across the three experimental blocks, we found that dependence on previous target direction was already reduced compared to reports without feedback by the end of the first block, with both auditory feedback (model-based: *a* = 1.209°, 95% CI: [-1.5421°, 2.2492°], *p* = .578; model-free: *M =* 1.0292°, *t*(9) = 1.1174, *p* = 0.293) and auditory + visual feedback (model-based: *a* = 2.088°, 95% CI: [-1.128°, 3.838°], *p* = .213; model-free: *M =* 2.796°, *t*(23) = 2.306, *p* = 0.031) compared to the no-feedback group, (model-based: *a* = 3.576°, 95% CI: [1.820°, 5.796°], *p* = .009; model-free: *M =* 4.202°, *t*(36) = 2.672, *p* = 0.011). The difference failed to reach significance by both measures for the auditory feedback and no feedback group comparison (model-based: lJ*a* = 2.555°, *p* = .101; model-free: Welch’s two-sample *t*(44.154) = 1.7408, *p =* .089; **Fig. 3c**). Likewise, the difference failed to reach significance by both measures for the auditory + visual feedback and no feedback group comparison (model-based: lJ*a* = 1.488°, *p* = .148; model-free: Welch’s two-sample *t*(58.928) = 0.7079, *p =* .48).

The model-based measure indicated no detectable serial dependence in blocks 2 and 3 with auditory feedback, whereas the model-free measure indicated a significant repulsive bias in block 2 (block 2: model-based: *a* = 0.0464°, 95% CI: [-7.1092°, 0.9545°], *p* = .459; model-free: *M* = −1.9897°, *t*(9) = 2.6201, *p* = 0.028; block 3: model-based: *a* = −0.4366°, 95% CI: [-2.7976°, 1.7943°], *p* = .951; model-free: *M* = −1.2362°, *t*(9) = 1.0245, *p* = 0.33). This significance repulsive bias was significantly different from the model-free estimate for both the no feedback and auditory + visual feedback conditions (*p* < .01 for both comparisons). With auditory + visual feedback, serial dependence was undetectable in blocks 2 and 3 by both measures (block 2: model-based: *a* = 1.241°, 95% CI: [-1.108°, 3.194°], *p* = .246; model-free: *M* = 1.421°, *t*(23) = 1.470, *p* = 0.155; block 3: model-based: *a* = 1.257°, 95% CI: [-1.310°, 4.064°], *p* = .512; model-free: *M* = 0.917°, *t*(23) = 1.102, *p* = 0.28).

Because our study design included additional manipulations, some of which were not relevant to the current analyses, we fit a linear mixed effects model to the model-free serial bias data that accounts for all manipulations. Main effects of (i) feedback (3 levels: auditory, auditory + visual, & without), (ii) current trial target contrast (3 levels: low, mid, & high), (iii) head-tracking condition (3 levels: fixed, active, & lagged), and (iv) block order (3 levels: first, second, third) were included in the analysis as fixed effects. Feedback and head-tracking condition were coded as categorical variables, and block order was coded as an ordinal variable. Additionally, the interaction between block and feedback was included. As random effects, the model included intercepts for subjects. Visual inspection of residual plots did not reveal any obvious deviations from homoscedasticity or normality. P-values were obtained by F-tests for each term in the linear mixed-effects model. The main effect of current trial target contrast was significant (F(1,627) = 25.321, p < .001). The main effect of feedback remained marginally significant when the feedback-block interaction was included in the model (F(2,627) = 2.844, p = .056), but was statistically significant when this term was left out (F(2,631) = 11.537, p < .001). There were no main effects of head-tracking condition (F(2,631) = 0.0996, p = .905) or block,(F(2,631) = 0.616, p = .541), and we did not find a block by feedback interaction (F(4,627) = 0.07716, p = .989). The failure to find a significant effect of block order is consistent with the notion that task feedback already affects performance in an observer’s first block of the task. It is worth noting that because there was variability in the effect of block due to the differential effects of head-tracking conditions on performance overall (see Fulvio & Rokers, 2017), specific effects of these factors are likely under-powered and obscured in this data set. This may also be an indirect source of the reduced effect of feedback when the interaction term was included in the model. A more targeted investigation of block order by feedback effects may be of interest for future research. We focus on one factor that directly impacts sensory uncertainty - target contrast - below.

### Current trial sensory uncertainty increases serial dependence

A sensory uncertainty account of serial dependence predicts that greater uncertainty in the estimate of the current target’s motion leads to greater reliance on recent experience. And, conversely, that greater uncertainty in the previous trial estimate should lead to a smaller impact of recent experience. Here, we tested both predictions by quantifying serial dependence as a function of our manipulation of target contrast.

We first measured serial dependence as a function of current trial target contrast. Increases in current trial target contrast were associated with reduced serial dependence on the previous trial’s target direction. In particular, serial dependence was abolished on high target contrast trials, consistent with a sensory uncertainty account (**Fig. 4a**). Without feedback, both reduced target contrast levels were associated with significant serial dependence according to both serial dependence measures (model-based: *a_high_* = 0.174°, 95% CI: [-2.631°, 0.927°], *p =* .265; *a_mid_* = 4.299°, 95% CI: [2.633°, 6.048°], *p <* .001; *a_low_* = 4.374°, 95% CI: [2.467°, 6.252°], *p =* .01; model-free: low: *M* = 4.269°, *t*(36) = 2.929, *p* = .006; mid: *M* = 3.467°, *t*(36) = 2.644, *p* = .012). With auditory + visual feedback, only the lowest target contrast level was associated with significant serial dependence according to both measures (model-based: *a_high_* = −0.976°, 95% CI: [-1.869°, 1.069°], *p =* .151; *a_mid_* = 0.987°, 95% CI: [-1.138°, 2.171°], *p =* .569; *a_low_* = 3.279°, 95% CI: [1.696°, 5.309°], *p <* .001; model-free: low: *M* = 2.41°, *t*(23) = 3.021, *p* = .006). With auditory feedback, none of the target contrast levels was associated with significant serial dependence according to both measures (*a_high_* = 1.438°, 95% CI: [-1.188°, 3.785°], *p =* .160; *a_mid_* = −1.485°, 95% CI: [-2.875°, 0.717°], *p =* .273; *a_low_* = 1.367°, 95% CI: [-2.605°, 2.498°], *p =* .132).

**Fig. 4.**
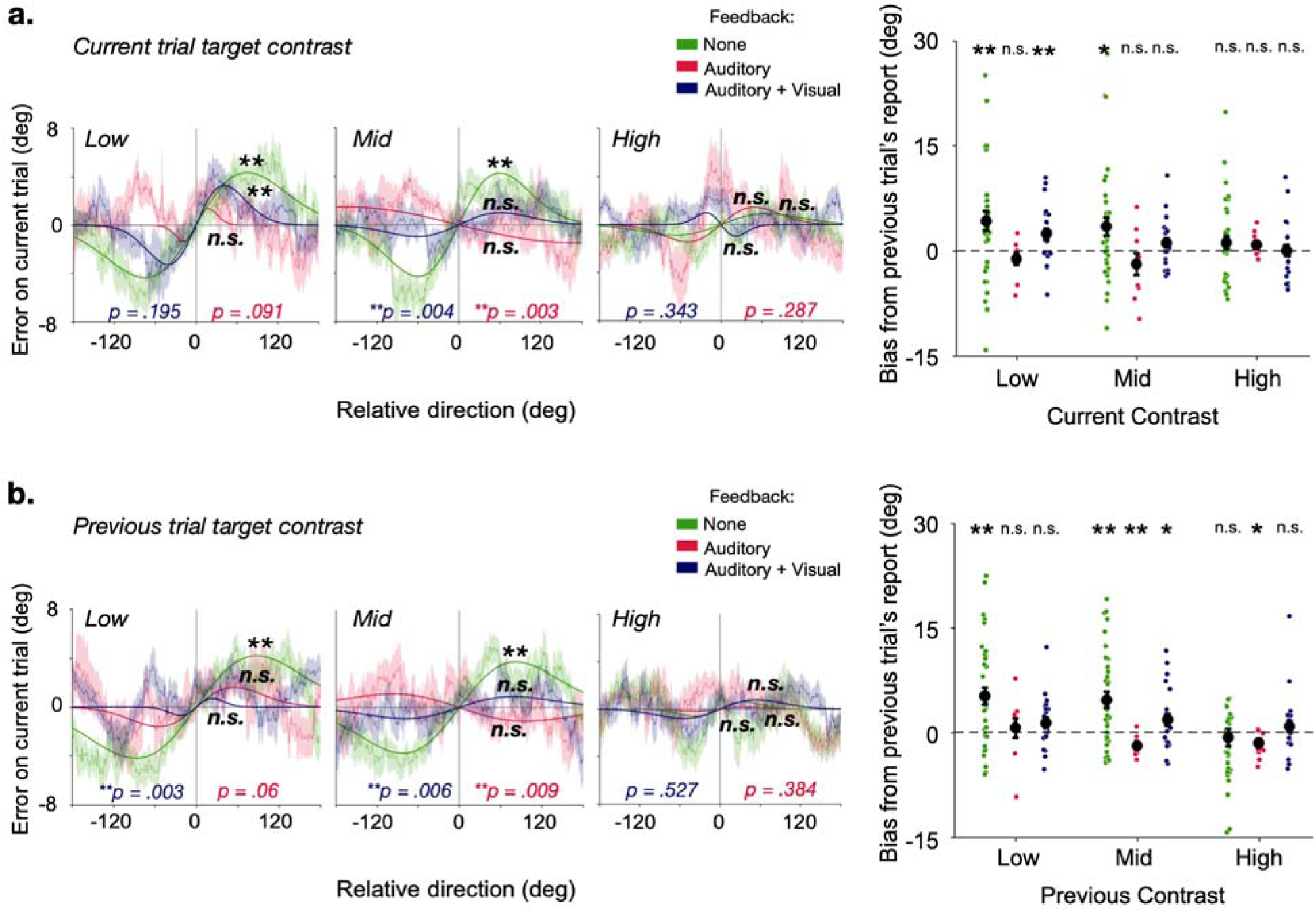
Serial dependence is modulated by sensory uncertainty due to target contrast. **(a.)** Left: Serial dependence curves and DoG fits to error data with high error trials omitted (i.e., on correct depth report trials following correct depth report trials), when no feedback was provided (green), when auditory feedback was provided (red), and when auditory + visual feedback was provided (blue), sorted according to the relative direction of the current trial’s target with respect to the previous trial’s target direction and split out by current trial target contrast. Shaded bands represent +/-1 *SEM*. **corresponds to bootstrapped *p*-values < .001; n.s. = non-significant; *p*-values at the bottom of each graph refer to the difference in amplitude parameter between no feedback and auditory feedback groups (red font) and between no feedback and auditory + visual feedback groups (blue font) derived from permutation testing. Right: Model-free serial bias with respect to the previous trial’s target direction for the three feedback conditions, split out by current trial target contrast. Circular symbols correspond to individual subject biases; black symbols correspond to group means. Error bars represent +/-1 *SEM*. **corresponds to *p*-values < .01; *corresponds to *p*-values < .05; n.s. = non-significant. **(b.)** Same formats as **(a.)** split according to previous trial target contrast.

Dependencies were significantly smaller when feedback was provided compared to when it was not provided for the mid target contrast level only (permutation test *p*-values ≤.0037 for both mid target contrast level comparisons; *p* ≥ .0913 for all other comparisons). Furthermore, analysis of the model-free serial bias measure revealed a significant effect of current trial target contrast for the no feedback group (*F*(1,109) = 6.147, *p* = .015) and for the auditory + visual feedback group (*F*(1,70) = 5.319, *p* = .024), but the effect failed to reach significance for the auditory feedback group (*F*(1,28) = 3.703, *p* = .065).

Interestingly, the relationship with the previous trial’s target contrast was less straight-forward (**Fig. 4b**). A sensory uncertainty based explanation might predict that greater target contrast on the previous trial would be associated with a more reliable estimate on that trial, which should have greater impact on the current trial’s estimate. With auditory + visual feedback, the model-based measure indicated no reliance on the previous target direction at any target contrast level (*a_previoushigh_* = 0.789°, 95% CI: [-1.225°, 1.558°], *p* = .586; *a_previousmid_*= 0.911°, 95% CI: [-1.417°, 1.909°], *p* = .792; *a_previouslow_*= 0.709°, 95% CI: [-1.379°, 1.663°], *p* = .902), whereas the model-free measure revealed a small positive bias on trials following mid-level target contrast trials only (*M* = 1.845°, *t*(23) = 2.0859, *p* = .048). The model-based measure also indicated no reliance on the previous target direction at any target contrast level for the auditory feedback group (*a_previoushigh_* = 0.173°, 95% CI: [-2.905°, 1.917°], *p* = .965; *a_previousmid_* = −1.124°, 95% CI: [-4.187°, 1.206°], *p* = .501; *a_previouslow_* = 1.610°, 95% CI: [-0.969°, 2.776°], *p* = .081), but the model-free measure revealed significant negative biases on trials following mid- and high-level target contrast trials (mid: *M* = −1.872°, *t*(9) = 4.218, *p* = .002; high: *M* = −1.525°, *t*(9) = −2.381, *p* = .041). However, in the no-feedback condition, reliance on the previous trial’s target direction was significant when the previous trial’s target contrast was mid or low, but not when it was high by both measures (model-based: *a_previoushigh_* = 0.604°, 95% CI: [-3.275°, 2.084°], *p* = .89; *a_previousmid_* = 3.769°, 95% CI: [2.268°, 5.283°], *p* < .001; *a_previouslow_*= 4.212°, 95% CI: [2.598°, 5.885°], *p* < .001; model-free: low: *M* = 5.243°, *t*(36) = 4.247, *p* < .001; mid: *M* = 4.703°, *t*(36) = 4.187, *p* < .001).

Dependencies were significantly smaller when auditory + visual feedback was provided compared to when it was not provided when the previous trial’s target contrast was low (permutation test *p*-value = .003) and trending in the same direction when auditory feedback was provided (permutation test *p*-value = .061). When the previous trial’s target contrast was at the mid-level, serial dependence was significantly smaller for the two groups in which feedback was provided compared to the no feedback condition (permutation test *p* ≤ .0088 for both comparisons). No differences were observed for the comparisons of serial dependence when the previous trial’s target contrast level was high (permutation test *p* ≥ .384 for both comparisons). This pattern of results suggests that the reliability of the current trial’s sensory estimate is a strong predictor of the degree of serial dependence in this task, but the reliability of the previous trial sensory reliability, less so (see also Gallagher & Benton, 2022).

In an additional analysis, we considered the *relative* contrast between the current and the previous trial and found that dependence on the previous trial’s target direction was only significant for the no feedback group when the current trial’s target contrast matched the previous trial’s target contrast according to both serial bias measures (*a* = 5.405°, 95% CI: [3.549°, 7.572°], *p <* .001; all other *p*-values ≥ .2448; model-free: *M* = 6.085°, *t*(36) = 4.127, *p* < .001; **Fig. S3**). In summary, these results suggest that sensory uncertainty as manipulated either through task feedback or stimulus contrast on the current trial, strongly modulates serial dependence.

## Discussion

Serial dependence – the impact of recent stimulus history on current perceptual reports (Fründ et al., 2014; Fischer & Whitney, 2014) – is a pervasive finding in many psychophysical domains. In the current study, we show that serial dependence occurs in psychophysical reports of 3D motion direction. Critically, however, we showed that serial dependence is not a given - reducing sensory uncertainty by increasing target contrast or providing performance feedback on a trial-to-trial basis was associated with near abolishment of serial dependence.

The target contrast-driven effects on serial dependence extend recent findings. In particular, the sensory reliability of the current stimulus was a stronger and more consistent predictor of serial dependence than the reliability of the previous stimulus. Whereas increases in current trial target contrast were associated with reductions in serial dependence for both feedback and no feedback groups, the effect of previous trial target contrast was less clear. We note that one potential limitation of our design was the use of above-detection threshold stimuli. Including lower contrast targets may have revealed serial dependence effects in the feedback conditions as well. Nevertheless, these results are consistent with other work showing that uncertainty in the current sensory estimate drives serial dependence more than previous trial uncertainty (e.g., Pascucci et al., 2019; Ceylan et al., 2021; Gallagher & Benton, 2022). Our results also support previous work showing that similarity between successive stimuli matters (Fritsche, 2016; Cicchini et al., 2018; Fritsche & de Lange, 2019; Lidström, 2019). By contrast, the feedback-driven attenuation of serial dependence is a novel finding. In fact, our results appear to contradict a recent report of *increased* attractive serial bias found in numerosity estimates with feedback (Fornaciai & Park, 2022). In that study, feedback was provided to participants in the first eight experimental blocks, followed by four blocks without feedback. Because serial dependence was smaller for the final four no feedback blocks compared to the four blocks with feedback, and on par with serial dependence observed in a previous study by the group in the absence of feedback (Fornaciai & Park, 2020), the interpretation was that feedback encouraged greater weight on the previous trial’s stimulus, especially when the feedback on the previous trial was ‘correct’. However, with the fixed block order design, it is not clear whether the effects were in fact feedback-based or merely experience-based. Indeed, in our study, we showed that as experience evolved in the task, serial dependence declined – even the no-feedback group’s performance exhibited hints of a decline in serial dependence over blocks. However, we emphasize that although feedback was associated with a general improvement in behavioral performance, including a reduction in response errors, those who learned to reduce response errors more over the course of the experiment did not necessarily reduce serial dependence more. Because the relationship between error variability and serial bias was not systematic, it does not seem that the effect of feedback on sensory reliability is the sole driver of the effect of feedback on serial dependence. Thus, taken together, both sets of results, those reported here combined with the recent work by Fornaciai & Park (2022) support the notion that serial dependence may originate at higher, post-perceptual levels, as also suggested by other results across the literature (Fritsche et al., 2017; Pascucci et al., 2019; Bae & Luck, 2020; Ceylan et al., 2021).

We note that in the auditory + visual feedback group, participants were presented again with the stimulus as well as their response during the feedback stage. Thus, participants in this group were provided with an opportunity to update their estimate of the motion direction and target endpoint on that trial. Nevertheless, this did not encourage continued dependence on the previous target direction beyond the first block. When the visual component was removed from the feedback for the auditory feedback group, serial dependence was, if anything, repulsive. The distinct nature of the feedback signals in these two conditions may nevertheless have contributed to the subtle differences in serial dependence between the two feedback groups. Taken together, our results raise the possibility that serial dependence is subject to strategic control, driven at least in part by error signals from feedback about the accuracy of one’s previous perceptual estimates.

In previous work, we interpreted improvements in accuracy of 3D motion-in-depth judgments with feedback as the result of the inexperienced participants learning to overcome “flatness” priors developed through extensive real-world use of 2D displays and to compel recruitment of the available depth cues that they were otherwise discounting (Fulvio & Rokers, 2017; Fulvio, Ji et al. 2020). Given that integration of multiple cues enhances reliability of visual perception (Chang et al., 2020; Hillis et al., 2004; Knill & Saunders, 2003; Murphy et al., 2013; Oruc et al., 2013; Preston et al., 2009; Rideaux & Welchman, 2018; Welchman et al., 2005), increasing the number of cues integrated in estimating the target’s motion direction may be one way in which feedback reduces serial dependence. (We note, however, that our design leaves open the possibility that other factors may also have accounted for the effects of feedback, given that adjustment responses are subject to some motor error and possible biases towards/away from certain directions. Future work employing other paradigms such as 2AFC tasks can help clarify the impact of feedback on sensory estimation in this task.) Nevertheless, the reduction in serial dependence in 3D motion reports observed in the current data set suggests that serial dependence may be a strategy employed specifically when the number or reliability of available sensory signals is low, supporting variable serial dependence across task conditions and trials within the same task context.

Feedback may also play a role in improving participants’ understanding of the task structure. Whereas real-world stimuli are often stable over various time scales such that taking information from the recent past into consideration when making current decisions could be adaptive (Fischer & Whitney, 2014; Kiyonaga et al., 2017; Braun et al., 2018), psychophysical stimuli, such as those in the current study, are often independent and (pseudo-)randomly presented by design. Serial dependence for unrelated stimuli is clearly suboptimal from an experimental point of view, but until participants are made aware of their mistakes, generalizing from experience provides a way of overcoming uncertainty. Indeed, we showed previously that participants in the no-feedback group exhibited a bias to report high contrast targets as approaching and reduced contrast targets as receding (Fulvio & Rokers, 2017), consistent with a dimmer-is-further-away response heuristic (Cooper & Norcia, 2014). In the current results, we show that serial dependence in the reports of the no-feedback group was significantly greater for reduced target contrast targets, thus supporting serial dependence as a behavioral strategy to overcome sensory uncertainty.

In conclusion, our results provide novel evidence supporting the notion that serial dependence can emerge as the result of a high level strategy employed to overcome sensory uncertainty. Our results are in line with the idea that serial dependence may reflect a flexible perceptual mechanism that is not limited to sensory components of external stimuli, but instead exploits all available information (Fornaciai & Park, 2022), including external feedback-based error signals.

### Citation diversity statement

To promote transparency surrounding citation practice (Dworkin et al. 2019; Zurn et al. 2020) and mitigate biases leading to under-citation of work led by women relative to other papers as demonstrated across several scientific domains (e.g., Dworkin et al. 2019; Maliniak et al. 2013; Caplar et al. 2017; Fulvio et al. 2021), we proactively aimed to include references that reflect the diversity of the field and quantified the gender breakdown of citations in this article according to the first names of the first and last authors using the Gender Citation Balance Index web tool (https://postlab.psych.wisc.edu/gcbialyzer/). This article contains 50.0% man/man, 12.5% man/woman, 26.6% woman/man and 10.9% woman/woman citations. For comparison, proportions estimated from articles in five prominent neuroscience journals (as reported in Dworkin et al. 2019) are 58.4% man/man, 9.4% man/woman, 25.5% woman/man and 6.7% woman/woman. Note that the estimates may not always reflect gender identity and do not account for intersex, non-binary, or transgender individuals.

## Funding

J.M.F. and B.R. were supported by funding provided by Facebook Reality and Google Daydream. The funders had no role in study design, data collection and analysis, decision to publish, or preparation of the manuscript.

## Notes

### Competing Interest Statement

The authors have declared no competing interest.

### Summary of Updates

Figure 2 revised; new table summarizing the results added; additional behavioral statistics included; minor revisions to descriptions of methods and statistical tests for clarity; minor revisions to Intro and Discussion to clarify interpretations and better position paper within the literature

