## Supplementary Materials for "Task feedback suggests a post-perceptual component to serial dependence"

**All-trial analysis**

In the main text, we report analyses of serial dependence in which high-error trials (i.e., trials in which the motion-in-depth was misreported) were omitted following the same analysis approach as previous studies (Bliss et al., 2017; Fritsche et al., 2017; Samaha et al., 2019). For completeness, we report the results of the primary analyses using the full set of trials here, henceforth, ‘all-trial analysis’.

We first replicated the analyses reported in **Figures 3a-b** that demonstrated a significantly reduced attractive serial dependence on the previous trial’s target direction with feedback (**see Fig. S1**). The attractive bias of the no feedback group did not emerge in the all-trial analysis by either model-based or model-free measures (model-based: *a* = 0.203°, *p* = .98; model-free: *M* = -1.3935°, *t*(36) = -1.685, *p* = .101). The model-based measure for the all-trial analysis indicated that auditory + visual feedback was associated with a significant *repulsive* bias (*a* = -1.43°, *p* = .02), whereas the model-free measure indicated a numerically repulsive bias that did not reach significance (*M* = -0.4145°, *t*(23) = -0.5331, *p* = .599). For the auditory feedback group, the model-based measure numerically indicated a repulsive bias as well, however, it failed to reach significance (*a* = -1.92°, *p* = .29), whereas the model-free measure did reveal a significant repulsive bias for the group that received auditory feedback (*M* = -2.873°, *t*(9) = -3.407, *p* = .008). Furthermore, neither measure of serial bias indicated significant differences between the no feedback and feedback group performances (model-based: *p* ≥ .0865 for both comparisons; model-free: *p* ≥ .0732 for both comparisons).

The all-trial analysis suggests a different nature of serial dependence in the 3D motion task than the high-error excluded analysis revealed in the main text. To reconcile the difference, we examined the relationship between presented and reported direction across trials, noting that in motion-in-depth misreport trials, the two are in opposition. Beginning with the group that did not receive feedback, the inclusion of the motion-in-depth misreport trials nulled any attractive serial biases observed when those trials were omitted. Inspection of participants’ responses more closely identified a tendency to report the same depth direction across trials independent of the presented direction. That is, we found that on 62.11% of trials participants who did not receive feedback reported the same depth direction on the current and previous trial, compared to effectively chance (50.31%) level of actual repeats in depth direction across successive trials. This difference was statistically significant (*t*(36) = 7.9896, *p* < .001). The apparent loss of serial dependence in the all-trial analysis for this group, then can be explained at least in part by this motion-in-depth report bias that obscures the results when high-error trials are included and the trials are sorted by relative *target direction* differences. The all-trials results thus also support the notion that the previous trial’s presented direction biases the current trial’s response.

Turning to the two groups that received feedback, the motion-in-depth response bias was significantly reduced for both groups compared to the no feedback group (*p* ≤ .0012 for both comparisons), but remained significant for the auditory + visual feedback group (53.77%; *t*(23) = 3.5304, *p* = .002; compared to the auditory feedback group (52.49%; *t*(9) = 1.945, *p* = .084)). More importantly, the feedback signal always indicated that the response was incorrect on motion-in-depth misreport trials, a factor that likely contributed to the tendency toward *repulsion* from the previous trial’s target direction.

Taken together, these findings support the removal of the high-error motion-in-depth misreport trials to aid the interpretation of the serial dependence analyses and support the notion that serial dependence is driven by factors beyond low-level sensory stimulus processing.

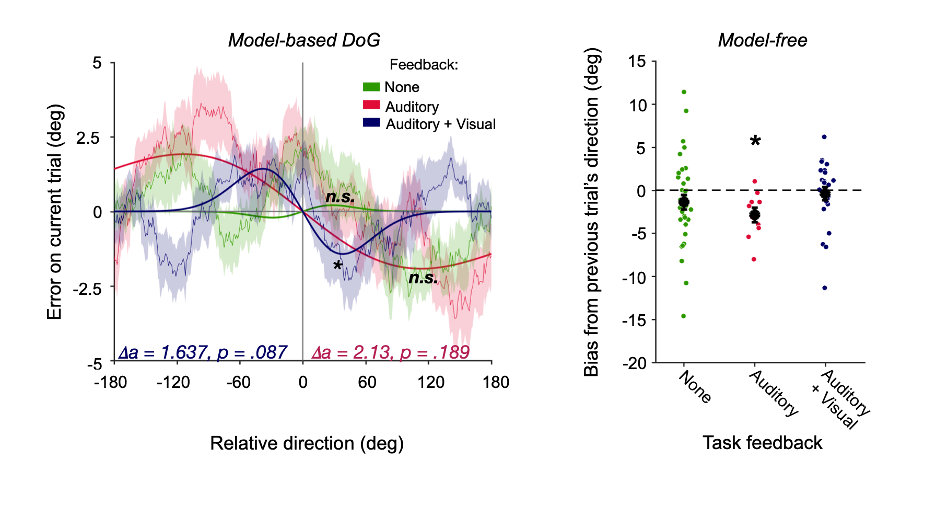

**Fig. S1**. **All-trial analysis of task feedback effects on serial dependence.** Left: Serial dependence curves and DoG fits to error data from all trials when no feedback was provided (green), when auditory feedback was provided (red), and when auditory + visual feedback was provided (blue), sorted according to the relative direction of the current trial’s target with respect to the previous trial’s target direction. Shaded bands represent +/-1 *SEM*. *corresponds to bootstrapped *p*-values < .05; n.s. = non-significant; 𝜟*a* refers to the difference in amplitude parameter between no feedback and auditory feedback groups (red font) and between no feedback and auditory + visual feedback groups (blue font) with corresponding *p*-values derived from permutation testing. Right: Model-free serial bias with respect to the previous trial’s target direction for the three feedback conditions. Circular symbols correspond to individual subject biases; black symbols correspond to group means. Error bars represent +/-1 *SEM*. *corresponds to *p*-value < .05.

For completeness, we also replicated the target contrast-based analyses reported in **Fig. 4** (see **Fig. S2**). When sorting trials according to the current trial’s target contrast, we found a *repulsive* bias according to the model-free measure only at the low target contrast level for the auditory feedback group (*t*(9) = -2.755, *p* = .022); otherwise, we found no significant serial dependence according to either measure for any of the three feedback groups for any of the three target contrast conditions (model-based: all bootstrapped *p*-values ≥ .071; model-free: *p* ≥ .071 for all remaining tests). Furthermore, no differences were observed between performance with and without feedback at any target contrast level (model-based: *p* ≥ .1435 for all permutation tests; model-free: *p* ≥ .0569 for all tests).

Importantly, these analyses are also impacted by response biases that obscure dependencies on previous target direction. For the group that did not receive feedback, we found significant repetition of reported depth direction over trials for all target contrast levels (*p* < .001 for all comparisons against base rate), with this tendency increasing as target contrast level decreased (*F*(2,108) = 22.758, *p* < .001). For the groups that did receive feedback, we did not find a significant main effect of target contrast level on the tendency to repeat reported depth directions (*p* ≥ .0875 for both tests). However, repetitions were significantly greater than expected according to the base rate for the low target contrast trials for both feedback groups (*p* ≤ .0057 for both tests). Additionally, for the auditory + visual feedback group, repetitions were significantly greater than expected for the high target contrast trials (*t*(23) = 2.7881, *p* = .0105) and trending for the mid target contrast trials at the Bonferroni-corrected alpha-level = .0167 (*t*(23) = 2.3433, *p* = .028).

When sorting trials according to the previous trial’s target contrast, we found significant *repulsive* biases when the previous target contrast level was low according to the model-based measure (*a* = -5.034°, *p* = .001) and when the previous target contrast level was mid and high according to the model-free measure (*p* ≤ .0302) for the auditory feedback group. The model-free measure also identified a repulsive bias when the previous target contrast level was high for the group that did not receive feedback (*M* = -2.67°, *t*(36) = -2.137, *p* = .0394). No other biases were identified (model-based: all bootstrapped *p*-values ≥ .2218; model-free: *p* ≥ .1641 for all tests). When the previous target contrast level was low, the bias was significantly more repulsive for the auditory feedback group compared to the group that did not receive feedback (𝛥*a* = 5.606°, *p* = .001); otherwise, no differences were observed between performance with and without feedback (model-based: *p* ≥ .114 for all remaining permutation tests; model-free: *p* ≥ .0936 for all tests).

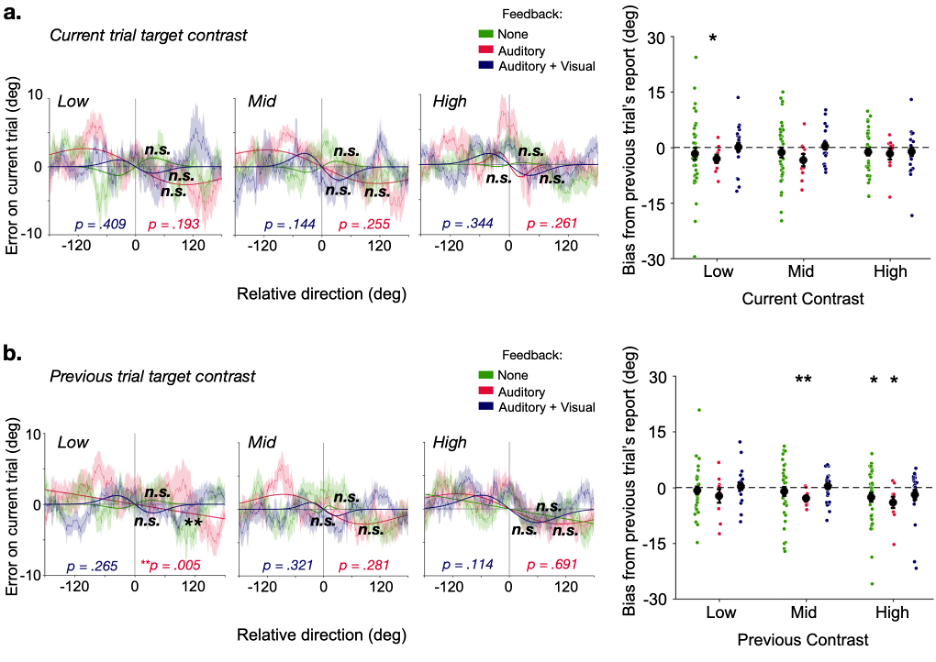

**Fig. S2.** **All-trial analysis of target contrast effects on serial dependence.** **(a.)** Left: Serial dependence curves and DoG fits to error data from all trials when no feedback was provided (green), when auditory feedback was provided (red), and when auditory + visual feedback was provided (blue), sorted according to the relative direction of the current trial’s target with respect to the previous trial’s target direction and split out by current trial target contrast. Shaded bands represent +/-1 *SEM*. **corresponds to bootstrapped *p*-values < .001; n.s. = non-significant; *p*-values at the bottom of each graph refer to the difference in amplitude parameter between no feedback and auditory feedback groups (red font) and between no feedback and auditory + visual feedback groups (blue font) derived from permutation testing. Right: Model-free serial bias with respect to the previous trial’s target direction for the three feedback conditions, split out by current trial target contrast. Circular symbols correspond to individual subject biases; black symbols correspond to group means. Error bars represent +/-1 *SEM*. *corresponds to *p*-values < .05 **(b.)** Same formats as **(a.)** split according to previous trial target contrast. **corresponds to *p*-values < .01.

**Relative target contrast analysis**

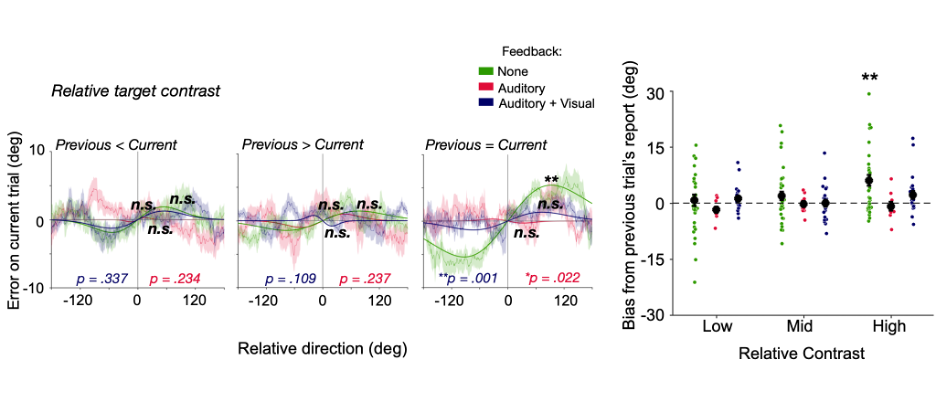

**Fig. S3.** **Serial dependence is modulated by relative target contrast.** **(a.)** Left: Serial dependence curves and DoG fits to error data with high error trials omitted (i.e., on correct depth report trials following correct depth report trials), when no feedback was provided (green), when auditory feedback was provided (red), and when auditory + visual feedback was provided (blue), sorted according to the relative direction of the current trial’s target with respect to the previous trial’s target direction and split out by current trial target contrast. Shaded bands represent +/-1 *SEM*. **corresponds to bootstrapped *p*-values < .001; n.s. = non-significant; *p*-values at the bottom of each graph refer to the difference in amplitude parameter between no feedback and auditory feedback groups (red font) and between no feedback and auditory + visual feedback groups (blue font) derived from permutation testing. Right: Model-free serial bias with respect to the previous trial’s target direction for the three feedback conditions, split out by current trial target contrast. Circular symbols correspond to individual subject biases; black symbols correspond to group means. Error bars represent +/-1 *SEM*. **corresponds to *p*-values < .01.
